# A newly-recognized population of residual neural crest cells in the adult leptomeninges is re-activated for vascular repair

**DOI:** 10.1101/2022.12.30.522316

**Authors:** Yoshihiko Nakamura, Takafumi Nakano, Ji Hyun Park, Masayoshi Tanaka, Wenlu Li, Elga Esposito, Bum Ju Ahn, Violeta Durán-Laforet, Rakhi Desai, Ikbal Sencan, Sava Sakadžić, Eng H. Lo, Evan Y. Snyder, Marcin Tabaka, Kazuhide Hayakawa

**Author notes:** **Correspondence:** Marcin Tabaka or Evan Y. Snyder or Kazuhide Hayakawa. **Author Contributions**, Performed experiments and/or analyzed data (YN, TN, JHP, MT^1^, EE, BJA, WL, VDL, RD, IS, ES); designed experiments (SS, MT^5^, ES, EHL, KH); wrote manuscript (TN, ES, MT^5^, EHL, KH); funding and support (EHL, KH).

## Abstract

The neural crest (NC) is a transient structure in vertebrate embryogenesis comprising highly migratory multipotent stem cells that give rise to a diverse array of cell types in organs throughout the body, including initiating neurovascular patterning. It is assumed that neural crest stem cells (NCSCs) disappear after development. Unexpectedly, using single-nucleus RNA-sequencing, we discovered residual quiescent NCSCs in the adult mouse meninges which are activated by injury and contribute to the brain’s homeostatic response. RNA velocity, pathway, and transcription factor analyses in a murine stroke model (combined with in vivo imaging) show that these adult NCSCs migrate towards the perivascular spaces of the infarct and undergo a perivascular stromal cell transition that is regulated by Ptp1b, Ghr, and Stat3. Loss- and gain-of-function experiments show that these “vestigial” NCSCs are required for restoring vascular endothelial barrier function via β-catenin and Stat3 signaling. These findings suggest that, in the adult, an unexpected reservoir of cells -- once pivotal to embryogenesis and vascular morphogenesis -- are re-invoked for neurovascular repair.

## Introduction

The neural crest (NC) is a transient structure in early vertebrate embryogenesis located on the dorsal neural primordium that consists of highly migratory, extremely multipotent stem cells that give rise to a diverse array of organs and cell types throughout the body (e.g., the peripheral, autonomic, and enteric nervous systems; bones of the face and cranium as well as the meninges; the great vessels of the heart; endocrine organs such as the adrenals and parathyroids; melanocytes; vascular smooth muscle) (Batarfi, 2017; Etchevers et al., 2001; Simoes-Costa and Bronner, 2016; Soldatov et al., 2019; Weigele and Bohnsack, 2020). They have also been found recently to be pivotal to establishing initial neurovascular patterning in the mammalian body plan shortly after gastrulation, even in the central nervous system (CNS) (Acevedo et al., 2015). It has been assumed that such neural crest stem cells (NCSCs) essentially disappear after development, perhaps giving rise to committed lineage-specific precursors/progenitors that reside in their respective organs (Dupin and Sommer, 2012) though no longer neural crest per se. Certainly the question remains open pending rigorous molecular characterization, a contribution to which we make here.

We became intrigued with an old observation that, in the adult CNS, a structure that originally derived from the neural crest and is conventionally viewed as fairly “inert” throughout life, serving principally as a “covering for the brain” – the leptomeninges -- may be “energized” by CNS pathology. Indeed, there has been a suggestion that meningeal involvement may be more than simply being a passive bystander to and recipient of the fallout from CNS injury; cells from the meninges have been observed to migrate into injured cortex and spinal cord and even generate neurons (Bifari et al., 2017; Decimo et al., 2011; Decimo et al., 2012; Siegenthaler et al., 2009). But such CNS progenitors are not NC derivatives; they are neural tube derivatives that happen to become sited at the brain’s surface (Decimo et al., 2020; Nakagomi et al., 2011). Knowing that the meninges themselves penetrate CNS parenchyma because they form the perivascular space around parenchymal blood vessels (Decimo et al., 2012; Hallmann et al., 2005) and are performing a “neural crest” role, we wondered whether there could, indeed – unexpectedly – be NCSCs “left over” from embryogenesis, even in the adult, that continue, to perform a normal neural crest role when “called upon”. It is, of course, known that certain adult neoplasias have a neural crest origin (e.g., meningioma, melanoma, pheochromacytoma); “embryonic rests”, so to speak. However, we wondered whether *normal* NCSCs may have been retained “<u>vestigially</u>” in the adult to continue performing normal neural crest-related homeostatic functions, for example, restoring vascular integrity after injury.

## Results

### Single-nucleus (sn) RNA-seq revealed gene signature of neural crest stem cells in the adult leptomeninges

To investigate this possibility, we performed deep characterization of the changes in the adult leptomeninges in response to a prototypical vascular-based adult CNS injury, i.e. stroke. First, we performed single-nucleus (sn) RNA-seq in male C57BL/6J mice (12-14 wks) subjected to distal middle cerebral artery occlusion (dMCAO) (3, 7, 14 and 28 days post-stroke) or sham-operated mice (3 days post) (**Fig. 1A**). The meninges are composed of 3 layers, arranged as follows from the brain outward to the skull: the pia mater, the arachnoid mater, and the dura mater. For these snRNA-seq analyses, we separated the brain – along with the adherent pia and a part of the arachnoid -- from the skull. We then dissected out the ipsilateral cerebral hemisphere. Therefore, the subsets identified in this experiment come from ipsilateral leptomeninges but not dura matter.

**Figure 1:**
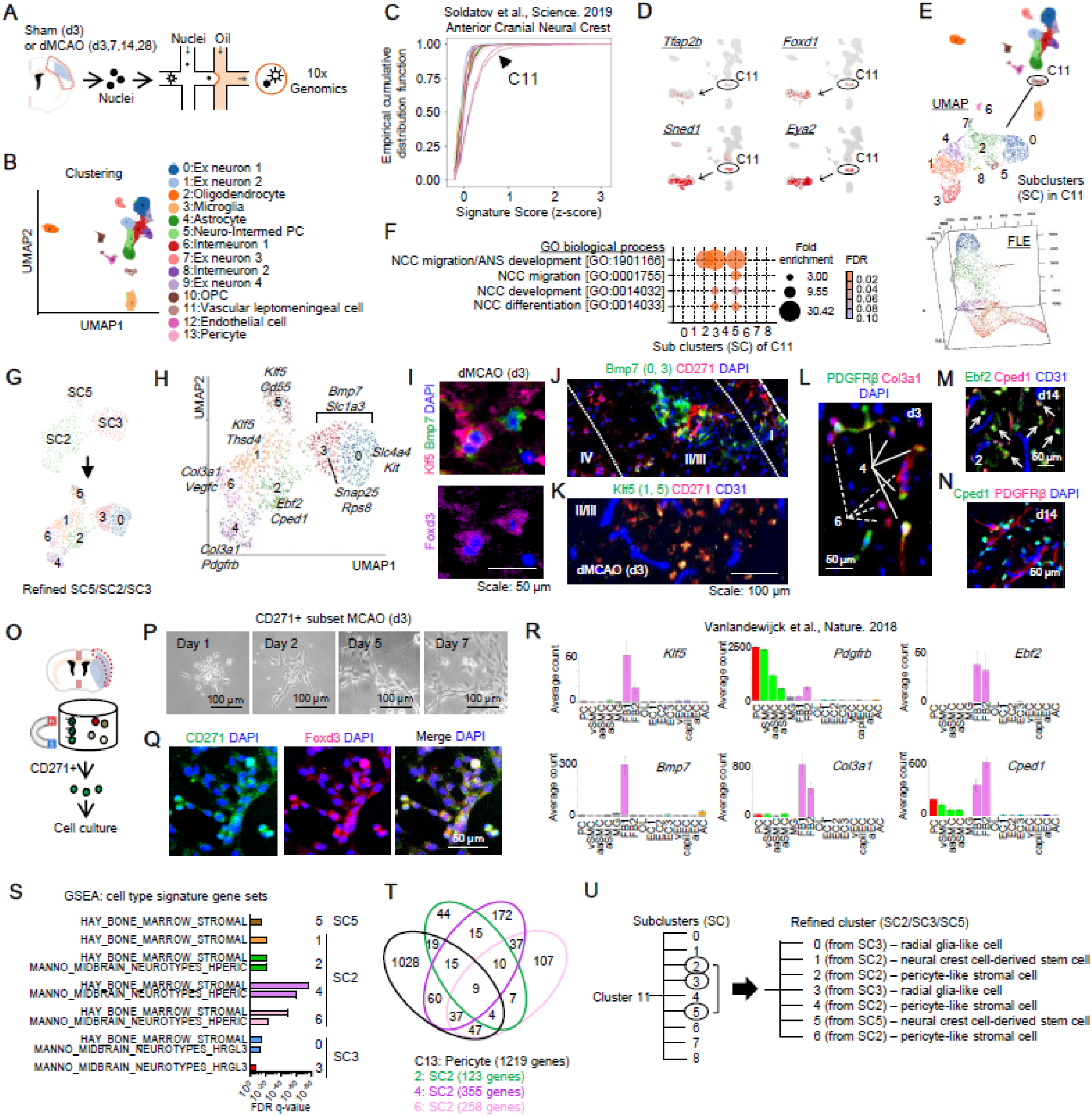
Identification of a repository of neural crest stem cells (NCSCs) in the adult leptomeninges as identified using single-nucleus (sn) RNA-seq. **A.** Single nuclei were extracted from the cortices of mice subjected to dMCAO 3 (n=3), 7 (n=4), 14 (n=3), and 28 (n=4) days before compared to control nuclei isolated from sham-operated age-matched mice (3 days post-operation, n=3). **B.** Unsupervised clustering of 107,020 nuclei, ranging from 16,411 to 29,570 nuclei across **the** above-mentioned time points using the R package, identified 14 clusters. PC: progenitor cells. **C.** Signature score analysis (z-score), using a *Wnt1*-associated early neural crest gene signature, revealed a subpopulation in *<u>Cluster 11</u>* as scoring highest (**arrowhead**). **D.** Cranial neural crest regulatory genes such as *Tfap2b, Foxd1, Sned1* and *Eya2* were found in *Cluster 11*. **E.** Upregulated “differential gene expression (DEG)” profiling in Cluster 11 in UMAP and FLE plot. **F.** “Gene Ontology (GO)” enrichment analysis identified, within *<u>Custer 11</u>*, subclusters *<u>SC2</u>*, *<u>SC3</u>*, and *<u>SC5</u>* as being involved in the regulation of neural crest migration. **G.** Subclusters *<u>SC2</u>*, *<u>SC3</u>*, and *<u>SC5</u>* were more finely resolved into Subsets 0-to-6 as follows: SC2 (Subsets 1, 2, 4, and 6), SC3 (Subsets 0 and 3), SC5 (Subset 5 alone). **H.** Upregulated genes in each Cluster. **I.** Immunohistochemical analysis showed that Klf5+ cells (Subsets 1 and 5) and Bmp7+ cells (Subsets 0 and 3) co-localized with Foxd3 (a transcriptional repressor involved in early NC commitment in the epiblast). Klf5 is a transcription factor critical for vascular survival and function. BMP7, a member of the TGF-β superfamily, plays a key role in bone homeostasis (transformation of mesenchymal cells into bone and cartilage) and is a well-accepted neural crest marker. PDGFRβ plays a critical role in the cranial neural crest contribution to craniofacial development, while Col3a1 is involved in the signaling necessary for neural crest cell migration. Ebf2 is a transcription factor that is expressed in E8.5 mesenchyme which is composed of neural crest and paraxial mesoderm. Cped1 is classified as a neural crest-associated gene though its precise function remains unknown. **J.** CD271+ cells were found in cortical layers I-to-IV ipsilateral to the dMCAO 3 days post-infarction. A subset of Bmp7+ cells (subsets 0 and 3) co-localized with CD271. **K.** Klf5+ cells were abundantly co-stained with CD271 in cortical layer II/III. **L-N.** The subsets of SC2 were found in the ipsilateral cortical layers II/III and IV, as confirmed by immunohistochemistry, as follows: Subset 2: Ebf2+/Cped1+ cells; Subset 4: PDGFRβ+/Col3a1+ cells; Subset 6: PDGFRβ-/Col3a1+ cells. **O.** CD271+ cells were isolated from leptomeninges-enriched tissue 3 days following transient focal cerebral ischemia for 60 min. **P.** Isolated cells were expanded in DMEM/F12 media plus 1% N2, 2% B27, FGF2 (100 ng/ml), and IGF1 (100 ng/ml). **Q.** Immunocytochemistry confirmed that isolated cells expressed CD271 and Foxd3. **R.** Representative genes in the Subsets of SC2, SC3, and SC5 were highly expressed in vascular cells (http://betsholtzlab.org/VascularSingleCells/database.html). **S.** Not unexpectedly, GSEA analysis also showed constituents of the meninges that were not neural crest-derived: Subset 4 was enriched for genes associated with bone marrow stroma and pericytes; Subsets 0 and 3 contained genes associated with neural tube-derived radial glia-like cells. **T.** Venn diagram of genes expressed in pericyte cluster C13 and in Subsets 2, 4, and 6. **U.** <u>Cluster</u> tree. Cluster 11 was divided into 9 distinct “subclusters”. Neural crest cell genes were enriched in Subclusters SC2, SC3, and SC5. These Subclusters were further divided into 7 distinct Subsets based on upregulated DEGs. **Abbreviations:** PC - Pericytes; SMC - Smooth muscle cells; MG - Microglia; FB - Vascular fibroblast-like cells; OL - Oligodendrocytes; EC - Endothelial cells; AC - Astrocytes; v - venous; capil - capillary; a - arterial; aa - arteriolar; 1,2,3-subtypes.

Unsupervised clustering of a total of 107,020 nuclei, ranging from 16,411 to 29,570 nuclei across the time points using the R package, identified 14 distinct clusters based on upregulated “differentially expressed genes (DEGs)” (**Fig. 1B, Fig. S1, Table 1, Table 2**). To find a “neural crest-derived cluster”, we examined the expression level of RNA signatures of *Wnt1* -based early neural crest as published in Soldatov et al (Soldatov et al., 2019). We found that leptomeningeal cell cluster 11 (C11) had the highest z-scores with neural crest gene signatures (**Fig. 1C**). Moreover, general cranial neural crest regulators including *Tfap2b, Foxd1, Sned1* and *Eya2* (Barque et al., 2021; Simoes-Costa and Bronner, 2016; Weigele and Bohnsack, 2020; Williams et al., 2019) were specifically upregulated in C11 (**Fig. 1D**). Next, C11 was divided into nine subclusters (SC0-8) based on DEGs (**Fig. 1E**). Gene ontology (GO) enrichment analysis revealed that SC2, SC3 and SC5 were enriched with genes regulating neural crest cell migration (**Fig. 1F**). Interestingly, these subclusters shared GO enrichment in neural crest-related mesenchymal cell differentiation and proliferation (**Fig. S2**), suggesting that SC2, SC3 and SC5 may be related to the formation of the meninges during embryogenesis. To gain a better understanding of the constitution of this neural crest derived population, we refined SC2, SC3 and SC5 subclusters using a shared nearest neighbor (SNN) modularity optimization-based clustering algorithm. This process divided SC2 into four distinct subsets (subsets 1, 2, 4, 6) and SC3 into two distinct subsets (subsets 0, 3), while SC5 was maintained as a single cluster (**Fig. 1G**).

Based on these refined DEGs (**Fig. 1H, Fig. S3, Table 3**), we next performed immunohistology to visualize the localization of these responses in the stroke-injured brains. NCSC markers such as Foxd3 (a transcriptional repressor involved in early NC commitment in the epiblast) and CD271 (p75 or the nerve growth factor [NGF] receptor [NGFR], and the most specific to NC (Lee et al., 2007)) were highly expressed in refined subsets 1 and 5, which also highly expressed Klf5 (a transcription factor that mediates the link between mitochondrial dynamics and NC-derived vascular smooth muscle cell [VSMC] survival and was highly expressed in cortical layer I-to-IV ipsilateral to the infarct at d3 poststroke) (**Fig. 1I-K**). Other refined SC2 subclusters such as PDGFRβ+/Col3a1+ cells (subset 4 from SC2) and PDGFRβ-/Col3a1+ cells (subset 6 from SC2) were identified in close proximity to perivascular areas (**Fig. 1L**), while Ebf2+/Cped1+ cells (subset 2 from SC2) localized to both perivascular and parenchymal regions in cortical layers II-III and IV by immunohistochemistry (**Fig. 1M-N**).

The presence of NCSCs was further confirmed by isolating CD271+ subsets from ipsilateral leptomeninges-enriched tissue via magnetic-activated cell sorting (MACS) (**Fig. 1O**) which is a well-accepted approach for isolating and culturing neural crest cells (Lee et al., 2007). Notably, CD271+ cells isolated at 3 day post-stroke were expandable in vitro (**Fig. 1P**) and consistently expressed NCSC markers (**Fig. 1Q**).

To further assess these NCSCs, we turned to a single-cell RNA-seq database (Vanlandewijck et al., 2018) to look for neural crest-associated genes. Our snRNA-seq data clearly showed expression of numerous neural crest-associated genes that are critical for vascular function, including *Klf5, Bmp7, Pdgfrb, Col3a1, Ebf2* and *Cped1* (**Fig. 1R**). Klf5’s role was described above. BMP7, a member of the TGF-β superfamily, plays a key role in bone homeostasis (transformation of mesenchymal cells into bone and cartilage) and is a well-accepted NC marker. PDGFRβ plays a critical role in the cranial neural crest’s contribution to craniofacial development, while Col3a1 is involved in signaling neural crest migration. Ebf2 is a transcription factor that is expressed in E8.5 mesenchyme which is composed of neural crest and paraxial mesoderm. Cped1 is assigned to neural crest-associated gene clusters although its function remains unknown. Refined subsets 1, 2, 4, 5, and 6 in Gene Set Enrichment Analysis (GSEA) contained enriched genes of bone marrow stromal cells. As per prior reports, SC3 subsets did contain enriched genes of radial glia-like cells that would have migrated there from the neural tube (*2-5*), but would not have been neural crest derivatives (**Fig. 1S**). Intriguingly, subsets 2, 4, and 6 also showed gene enrichment in pericytes (**Fig. 1S**), a cell type which, together with NC-derived VSMCs, is critical for vascular integrity. Therefore, we compared the NCSC cluster with the pericyte cluster C13. A Venn diagram shows that subsets 2, 4, and 6 shared 38.2% (47 out of 123 genes), 34% (121 out of 355 genes) and 37.6% (97 out of 258 genes) with the pericyte cluster, respectively (**Fig. 1T**).

Altogether, gene profiling and immunohistology demonstrated **(Fig. 1U)** that neural crest derivatives and residual NCSCs were most prominent in adult leptomeningeal Cluster 11, which, upon further refinement, could be localized to Subclusters (SCs) 2, 3, and 5 (SC2, SC3, SC5), and even more precisely to Subset 1 from SC2 and Subset 5 from SC5. Importantly, Subsets 2, 4, and 6 from SC2 were pericyte-like stromal cells, derived from mesoderm and the functional partners of neural crest in the vasculogenic process. Subsets 0 and 3 SC3, on the other hand, were not vasculogenic, but rather neural tube-derived neurogenic-radial glia-like cells (characterized by *Slc1a3* [also known as GLAST] expression (*Bifari et al., 2017)*).

### Leptomeningeal NCSCs transit to non-neural stromal cells through β-catenin and Stat3 pathways

Next, we assessed cell transition with the RNA velocity method (La Manno et al., 2018) which uses spliced and unspliced forms of RNA transcripts to infer potential cellular trajectories. The bam files from the 10x Genomics Cell Ranger pipeline were processed using Velocyto package (La Manno et al., 2018) to determine RNA expression matrices. The averaged percentage of spliced or unspliced RNA was 31.7% or 68.3% (sham), 25.6% or 74.3% (d3), 27.8% or 72.3% (d7), 31.3% or 68.6% (d14), and 32% or 68% (d28). Notably, subset 5 (from SC 5) committed transition to each subset with the following transition confidence: 0 (0.03), 1 (0.18), 2 (0.2), 3 (0.016), 6 (0.15), while refined Cluster 1 further transited to Cluster 4 (0.31) (**Fig. 2A**). PAGA graph abstraction (Wolf et al., 2019) with velocity-inferred directionality determined transitions between cell clusters; then PAGA velocity-graphs were visualized on the UMAP (Uniform Manifold Approximation and Projection) embedding (**Fig. 2B)**. Time course plots normalized by total cell numbers of Cluster 11 revealed that all SC2 subsets (1, 2, 4, 6) were stroke-induced populations while SC3 and SC5 were present in sham-operated controls (**Fig. 2C**).

**Figure 2:**
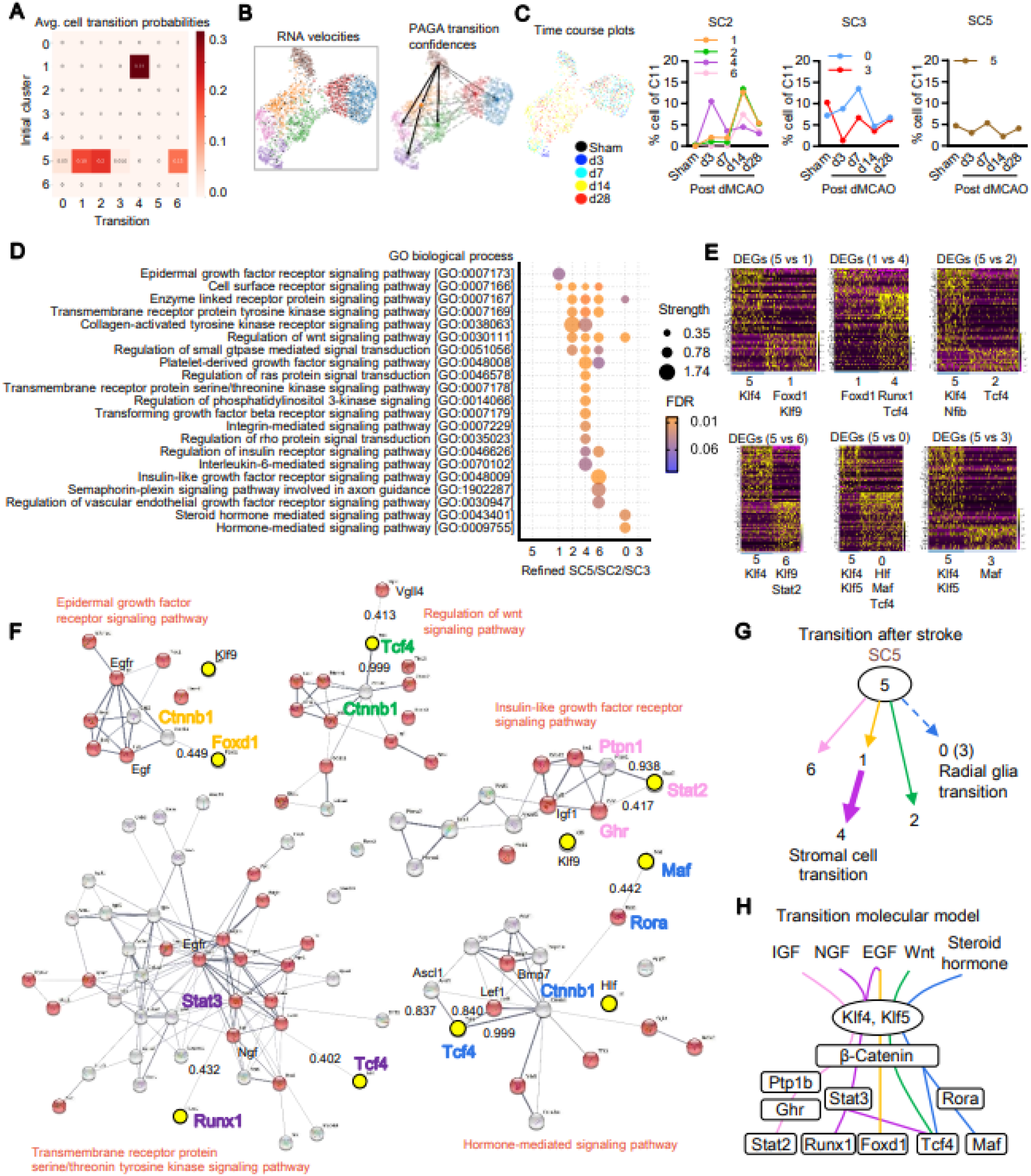
scVero analysis reveals trajectory of leptomeningeal NCSCs to non-neural neural crest-derived stromal cells through β-catenin and Stat3 pathways. **A.** Averaged cell transition probabilities revealed higher probabilities of SC5 to SC2 transition than SC5 to SC3. The transition of SC5 to SC1 to SC4 showed highest transition probability. **B.** scVelo indicated the transition of SC5 (Subset 5) toward SC2 (Subsets 1, 2, 4, and 6) but very low transition of SC5 (Subset 5) toward SC3 (Subsets 0 and 3) (See **Fig. 1u** for definition of the Subsets). **C.** Umap plots colored by time course post-dMCAO. SC2’s subsets (1, 2, 4, and 6) appeared after stroke while SC3’s subsets (0 and 3) and SC5’s only subset (Subset 5) were present in control. **D.** Gene Ontology (GO) enrichment analysis in refined subsets 0-6 demonstrated gene enrichment in each subset upregulating various signaling and regulatory pathways. **E.** <u>“Differentially expressed gene (DEG)”</u> analysis identified differential expression of transcription factors between transition subsets. **F.** Protein-protein interaction (PPI) network analysis along with uniquely upregulated signaling pathways and transcription factor(s) in each subset. **G.** SC5 subset transition to other subsets after stroke. Thicker arrow indicates higher transition probability while broken arrow implicates low transition probability. **H.** Transition molecular model. SC5’s Subset 5 transits to other subsets through key regulators such as upstream Ctnnb1 (β-catenin) followed by Ptpn1 (Ptp1b), Ghr, Stat3 or Rora and key transcription factors such as Stat2, Runx1, Foxd1, Tcf4, or Maf after stroke.

Taken together, these analyses suggest that, after stroke, neural-crest-derived meningeal cells in SC5 migrate and show the potential of mesenchymal non-neural progeny that has been described in neural crest cells isolated from ES cells (Lee et al., 2007). To assess the molecular pathway underlying this SC5 trajectories, we searched potential cell signaling pathways in GO biological processes based on a “false discovery rate (FDR)” <0.05 calculated by the Benjamini-Hochberg procedure. Subset 1 (from SC2) specifically upregulated *EGF receptor signaling pathway*. Subset 4 (from SC2) upregulated many signaling and regulatory pathways including *ras protein transduction, TGF-β signaling* and *integrin-mediated pathways*, and Subset 6 (from SC2) upregulated *insulin-like growth factor receptor signaling, semaphorin-plexin signaling*, and *VEGF receptor signaling* pathways. Moreover, Subset 0 (from SC3) uniquely increased genes regulating hormone-mediated signaling pathway (**Fig. 2D**). We also determined transcription factors differentially expressed in each transition partner (**Fig. 2E**) and subsequently performed “protein-protein interaction (PPI)” network analysis between signaling pathways and transcription factors to identify key molecules that potentially mediate each transition signal. Intriguingly, β-catenin (*Ctnnb1*) was found in the hub position of three key pathways potentially regulating fate decisions in neural crest cells (Brundage et al., 2014; Garcez et al., 2009; Hari et al., 2002): the *EGF receptor signaling* network along with *Foxd1*, regulation of *wnt signaling* network along with *Tcf4*, or *hormone-mediated signaling network* along with *Tcf4* or *Rora/Maf* interaction (**Fig. 2F**). Moreover, *Stat3* was the hub protein that interacts with both *Runx1* and *Tcf4* in the transmembrane receptor protein serine/threonine tyrosine kinase signaling network (**Fig. 2F**).

Taken together, RNA velocity and transition probabilities demonstrated that Subset 5 (from SC5) may change RNA transcripts that are similar to mesenchymal non-neural neural crest-derived stromal cells, especially Subset 4 (from SC2), through the state of Subset 1 (from SC2) with the *highest* probability via various regulatory mechanisms such as *Ctnnb1, Ptp1b, Ghr*, and *Stat3* (**Fig. 2G-H, Fig. S4**).

### Leptomeningeal NCSCs migrate to perivascular regions in the infarcted adult mouse cerebrum

So far, snRNA-seq identified a SC5 subset with gene enrichment in neural crest cells coming from a leptomeningeal cell cluster (C11) in adult mouse brain. However, snRNA-seq did not fully detect neural crest genes. Therefore, we sought to validate neural crest gene expression in cells that expressed the *nerve growth factor (NGF) receptor (NGFR*), i.e., *CD271 immunopositive cells*. In normal uninjured 12-13 week male C57BL/6J mice, CD271+ cells (demonstrated by immunohistochemistry) were predominantly located between the skull and the brain, within the cranial sutures and cranial bone marrow, but not in the cerebral cortex (**Fig. 3A**). We extracted leptomeninges-enriched tissue and isolated CD271+ cells by magnetic beads conjugated with a CD271 antibody (**Fig. 3B**). qPCR analysis confirmed that canonical neural crest genes such as *Tfap2, Foxd3*, and *Sox10* were highly expressed in CD271+ subset compared to bulk cells from the cerebral cortex (**Fig. 3C**).

**Figure 3:**
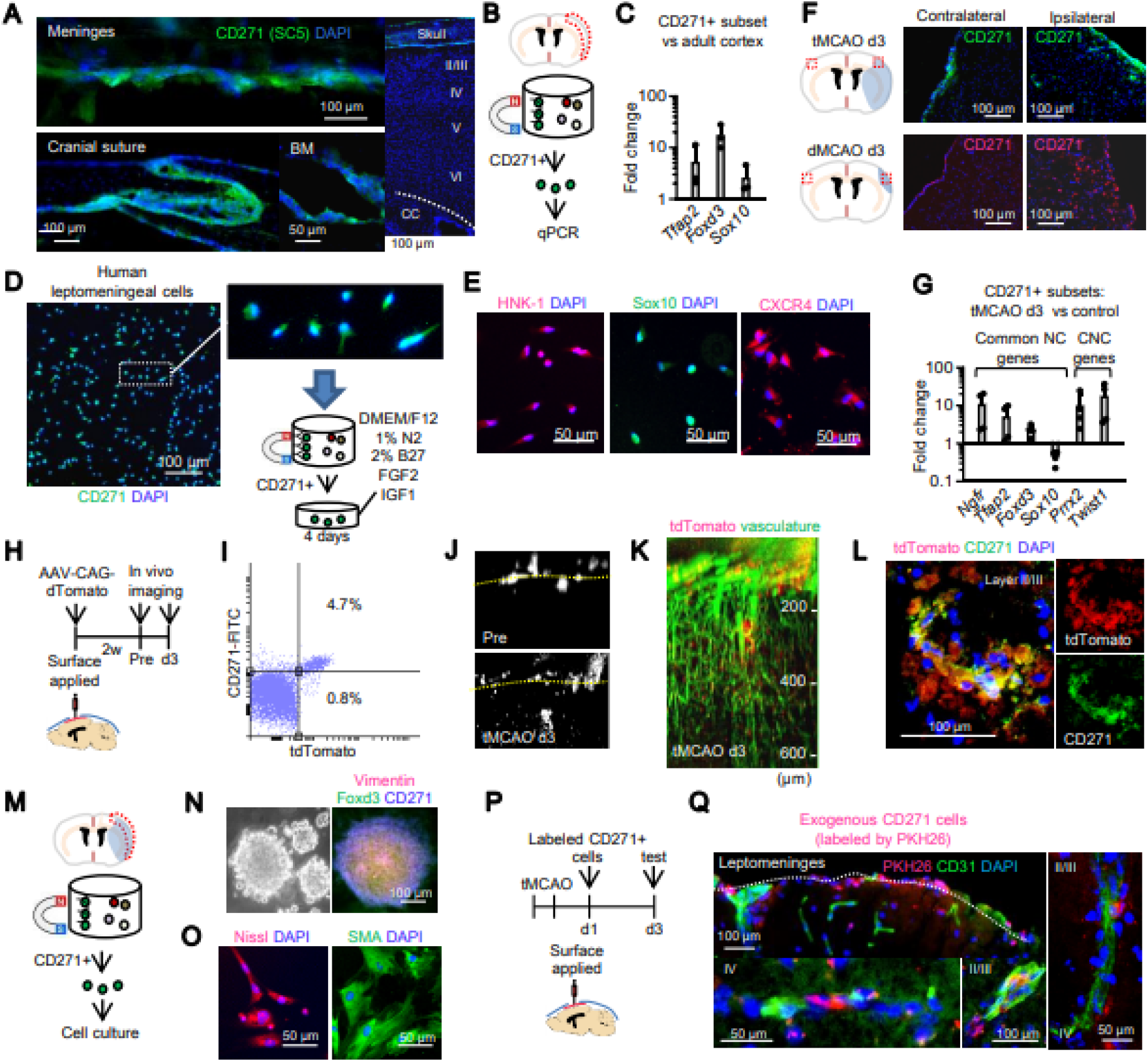
Migration and homing of leptomeningeal neural crest stem cells (NCSCs) to perivascular regions in the infarcted adult mouse cerebrum. **A.** In normal adult male C57BL/6J mice, CD271+ cells (Subset 5) were present in the meninges, cranial sutures, and skull bone marrow, but not in the cerebral cortex. **B-C.** CD271 cells isolated from leptomeninges-enriched tissue highly expressed *Tfap2, Foxd3, Sox10* compared to cells from the cortex. **D, E.** Not just the mouse meninges but also *human* leptomeninges contain CD271+ NSCSs. From primary human. **D.** A suspension of dissociated primary human leptomeninges were passed through magnetic beads conjugated with a CD271 antibody. The CD271+ cells were cultured for 4 days in DMEM/F12 plus 1% N2, 2% B27, FGF2 (100 ng/ml), and IGF1 (100 ng/ml). **E.** Immunocytochemistry confirmed that the purified human leptomeningeal CD271+ cells expressed the neural crest markers HNK-1, Sox10, and CXCR4. **F.** Immunostaining of the infarcted mouse brain showed that CD271+ cells were increased in the ipsilateral cortex at 3 days after transient (60 min) focal MCAO (tMCAO) or permanent distal MCAO (dMCAO). **G.** At day 3, after tMCAO, genes consistent with a neural crest identity and perivascular function, such as *Ngfr, Tfap2, Foxd3, Prrx2*, and *Twist1*, were upregulated in CD271+ cells while *Sox10* (a gene involved in neural crest’s melanocyte-yielding role) was decreased (Uninjured control mice, n=3; tMCAO mice, n=4). **H.** AAV-CAG-tdTomato (5×10^9^ viral genomes) was applied onto the ipsilateral leptomeninges via a cranial window in adult male C57BL/6J mice and two-photon imaging was conducted before and 3 days after tMCAO. **i.** Flow cytometry analysis confirmed that CD271+ cells (the label most specific for NC) were labeled with tdTomato. **I-K.** Some tdTomato+ cells migrated down to the ipsilateral cortex juxtaposed with brain penetrating vessels 3 days post-tMCAO. **L.** Immunostaining confirmed accumulation of CD271+ cells co-expressing tdTomato. **M.** CD271+ cells were isolated by MACS from the injured cortex after tMCAO. **N.** Leptomeninges-enriched fractions were extracted 3 days post-stroke. From this fraction, CD271+ cells were isolated by MACS. These CD271+ cells were capable of forming floating spheres that highly expressed vimentin and Foxd3. **O.** The multipotency of the CD271+ cells was confirmed by their ability to yield entirely different neural crest-derived lineages under different culture conditions. The addition of *nerve growth factor* (NGF; 50 ng/mL) and *brain-derived neurotrophic factor* (BDNF; 50 ng/mL) on 7 consecutive days (in the absence of serum) induced the emergence of peripheral or autonomic neurons (as detected by NeuroTrace-fluorescent dye obtained by Thermo Fisher Scientific) whereas TGF-β (5 ng/mL) and FBS (5%) in DMEM/F12 media for 7 days produced smooth muscle (as identified by the expression of smooth muscle actin [SMA]). **P.** Exogenous CD271+ cells were labeled ex vivo with the red fluorescent tracker PKH26 and transplanted onto the cortical surface at day 1 post-stroke. **Q.** At 3 days post-stroke, PKH26-labeled cells were in close proximity to blood vessels (recognized by their CD31 immunopositivity) in cortical layers II/III and IV. All data are shown as mean ± SD.

Importantly, by way of validation of our assessment in mouse, CD271+ cells were also found in *<u>human</u>* leptomeningeal cells and expressed cardinal neural crest markers (e.g., HNK-1, Sox10, and CXCR4) (**Fig. 3D-E**).

To assess the response to stroke, adult male C57BL/6J mice were subjected to transient middle cerebral artery occlusion (tMCAO) for 60 min. CD271+ cells were increased in the ipsilateral cortex by day 3 after tMCAO (**Fig. 3F**), suggesting that those cells responded to ischemic injury induced by both transient and permanent cerebral ischemia. After focal cerebral ischemia, neural crest-associated genes – such as *Ngfr* (CD271)*, Tfap2, Foxd3, Prrx2*, and *Twist1* – were increased, whereas *Sox10* expression was decreased in that CD271+ subset at day 3 post-stroke (**Fig. 3G**).

Because our snRNA-seq data suggested a capacity for both migration and transition predicted by the status of RNA transcripts, we next asked whether we could confirm movement of these CD271+ cells from leptomeninges into the ischemic brain. We performed two separate experiments.

First, we applied AAV-CAG-tdTomato directly onto the cortical surface through a cranial window to trace leptomeningeal cells at 2 weeks prior to 60 min tMCAO (**Fig. 3H**). Flow cytometry confirmed that CD271+ cells were labeled by tdTomato (**Fig. 3I**). In vivo two photon imaging showed that tdTomato+ cells migrated down to the ipsilateral cortex by 3 days post-stroke (**Fig. 3J-K**); immunostaining further confirmed the presence of CD271+ cells co-expressing tdTomato (**Fig. 3L**), suggesting the leptomeningeal origin of CD271+ cells that had accumulated in the damaged cortex.

Second, we used exogenous CD271+ cells as a tool to trace the migration pathway. After confirming sphere-forming ability and multipotency of the CD271+ cells (i.e., the ability to yield different neural crest derivatives under different inductive conditions) (**Fig. 3M-O**), they were prelabeled with the fluorescent tracer PKH26 *ex vivo* and placed (1×10^4^ cells) directly onto the ipsilateral cortical surface at 1 day after tMCAO (**Fig. 3P**). By 3 days post-stroke, PKH26+ cells were detected in close proximity to vasculature located in the ipsilateral cortical layers II-III and IV (**Fig. 3Q**), recapitulating our finding of the behavior of endogenous CD271+ cells post-stroke (**Fig. 1J**). Taken together, CD271+ cells within the leptomeninges appear to migrate into the perivascular space after stroke.

### Leptomeningeal NCSCs are required for restoring vascular endothelial barrier function via pleiotrophin-mediated phosphorylation of Akt and GSK-3β

To determine whether these putative endogenous NCSCs interacted with vascular endothelial cells (VECs) in the adult following stroke, as they did in the embryo during organogenesis, we next focused on the molecular crosstalk between these CD271+ cells and cerebral vasculature by performing a series of loss- and gain-of-function experiments. First, we looked for potential receptor-ligand interactions between SC5 and the brain VEC cluster (C12) in our snRNA-seq dataset. In VECs, *Cxcl12* (SDF-1α) showed the highest expression as a ligand to SC5 (**Fig. S5A**). Immunostaining and western blots confirmed that SDF-1α was increased in brain VECs and secreted into the cerebrospinal fluid (CSF) at 3 days after tMCAO (**Fig. S5B-C**). The SDF-1α receptor, CXCR4, is a well-known regulator for neural crest migration (Olesnicky Killian et al., 2009). Therefore, we next performed a loss-of-function experiment by blocking SDF-1α-CXCR4-mediated migration. To locally inhibit SDF-1α paracrine signaling, the selective receptor CXCR4 inhibitor, AMD3100, was directly applied onto the leptomeninges at day 2 post-stroke and outcomes were evaluated two days after the treatment (**Fig. 4A**). Treatment with AMD3100 significantly decreased the accumulation of cells expressing CD271 and Klf5 without affecting the number of F4/80+ macrophages or CD34 and Flk1 double positive vascular endothelial progenitors (**Fig. 4B-C**). Importantly, when acute migration of leptomeningeal CD271+ cells was inhibited, tight junction proteins Claudin-5 and ZO-1 were decreased (**Fig. 4D-E**) and, consistent with the fact that these regulators of vascular integrity were reduced, Evans blue extravasation through an impaired blood brain barrier (BBB) was significantly worsened (a functional measure) (**Fig. 4F-G**), although the infarct itself was not changed (**Fig. 4H-I**).

**Figure 4:**
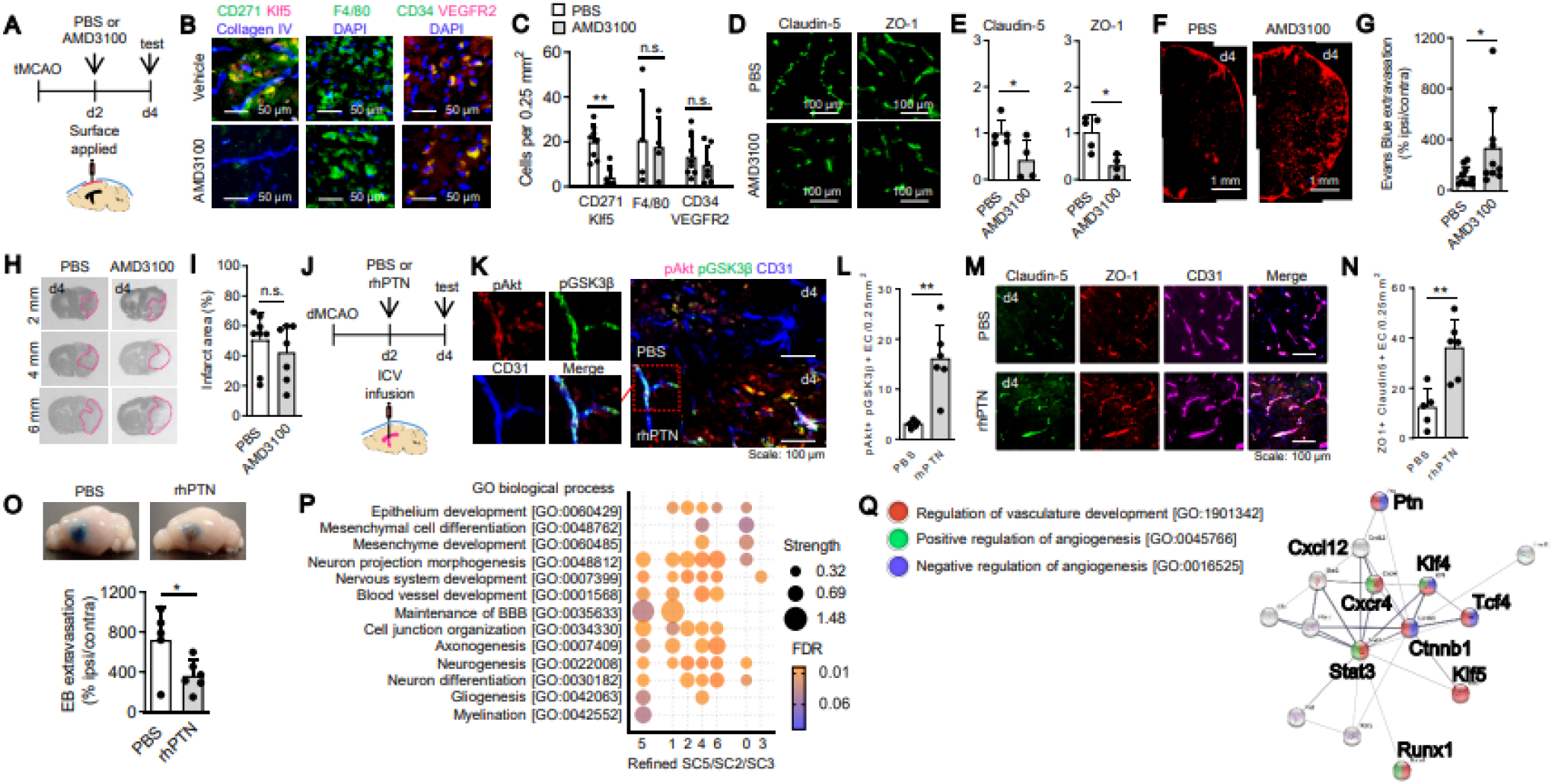
Restoration of the cerebral vascular endothelial barrier is mediated by NCSCs in the adult leptomeninges. **A.** Two days after focal cerebral ischemia, vehicle (PBS, 2 μL) or AMD3100 (a CXCR4-specific inhibitor, 10 μg/2 μL) was applied directly onto the cortical surface. **B-C.** Treatment with AMD3100 reduced the number of CD271/Klf5 double-positive cells per 0.25 mm^2^ in the peri-infarct cortex (PBS, n=8; AMD3100, n=7) but did not change the accumulation of macrophages (PBS, n=4; AMD3100, n=4) or vascular endothelial progenitors (PBS, n=8; AMD3100, n=7). **D-E.** AMD3100 decreased the expression of tight junction proteins Claudin-5 and ZO-1, as determined by immunocytochemistry (PBS, n=5; AMD3100, n=4). **F-G.** Functional integrity of the BBB was assessed by the degree to which it could prevent extravasation of Evans blue (1%, 100 μL) into the cerebrum following a system infusion (intravenously 30 min prior to cardiac perfusion and examination of the brain). AMD3100, which inhibited NCSC homing to the vasculature, but not changes in the vascular per se, significantly worsened Evans blue extravasation (PBS, n=10; AMD3100, n=10). **H-I.** Infarct area was quantified following Nissl staining. The infarct area [(contralateral hemisphere – non-infarct ipsilateral hemisphere) / contralateral hemisphere x100] was not significantly different in size when cell migration was blocked (Vehicle, n=8; AMD3100, n=7). **J.** Recombinant human Pleiotrophin (1 μg/5 μL) or vehicle (PBS, 5 μL) was infused intraventricularly at 2 days post-tMCAO. Evans blue extravasation, tight junction proteins (Claudin-5 and ZO-1), and pAkt/pGSK3β (Ser9) in brain vascular endothelial cells (VECs) were assessed at 4 days after stroke. **K-L.** Recombinant human Pleiotrophin significantly increased pAkt+ and pGSK3β (Ser9)+ VECs (CD31+) in peri-infarct areas (PBS, n=5; rhPTN, n=6). \*\**P*<0.01. **M-N.** Immunohistochemistry showed higher expressions of Claudin-5 and ZO-1 in CD31+ brain VECs in rhPTN-treated group compared to PBS-treated group. (PBS n=5, rhPTN n=6). \*\**P*<0.01. **O.** Treatment with recombinant PTN significantly attenuated Evans blue extravasation (PBS, n=5; rhPTN, n=6). \**P*<0.05. **P.** GO enrichment analysis demonstrated that Subset 5 and Subset 1 uniquely showed gene enrichment in maintenance of the BBB. **Q.** Protein-protein interaction (PPI) network analysis,_along with pleiotrophin (Ptn), SDF-1α (cxcl12), CXCR4, and genes from the transition model. Ptn interacts with Cttnb1 (β-catenin). Cttnb1 and Ptn commonly show GO enrichment involved in the regulation of vasculature development (which will insure BBB integrity) and negative regulation of angiogenesis (which makes for immature, leaky, inflammation-inducing vessels).

Next, we performed a gain-of-function experiment. Looking back into the snRNA-seq data again, a receptor-ligand interaction analysis between the endothelial cluster (C12) and SC5 identified *Ptn* (pleiotrophin) as the gene with the highest expression as a ligand connecting SC5 to C12 (**Fig. S5D-E**). To boost this endogenous pathway, we infused recombinant human pleiotrophin into the ventricles of mice at 2 days after dMCAO (**Fig. 4J**). Compared to vehicle controls, pleiotrophin treatment augmented Akt phosphorylation and serine-9-phosphorylation of GSK-3β in brain VECs (**Fig. 4K-L**), increased endothelial levels of claudin-5 and ZO-1 expression (**Fig. 4M-N**), and restored endothelial barrier function at d4 post dMCAO (again, as assessed by the prevention of Evan’s Blue extravasation from the systemic compartment into the brain) (**Fig. 4O**).

Finally, we asked whether these vascular repair signals could be detected in our original snRNA-seq data. GO biological process analysis of Subset 5 (from SC5) and Subset 1 (from SC2) (Klf5+, CD271+ cells) showed gene enrichment for maintenance of BBB integrity (**Fig. 4P**). PPI network analysis of the interactions among SDF-1α, CXCR4, pleiotrophin, and the molecules highlighted in the previously-described cell transition molecular model revealed that pleiotrophin interacted with Ctnnb1 (β-catenin) and Cxcl12 (SDF-1α) concomitant with gene enrichment for the positive regulation of vasculature development (with barrier integrity) and negative regulation of angiogenesis (**Fig. 4Q**), consistent with the knowledge that angiogenesis-induced new vessels have higher-than-normal permeability (i.e., they are immature, leaky, and allow for the ingress into the CNS of inflammatory cells from the systemic compartment) (Yang and Torbey, 2020). Furthermore, the SDF-1α receptor CXCR4 was strongly associated with Stat3 upregulation (**Fig. 4Q**) during cell transition to perivascular stromal Subset 4 (**Fig. 2G-H**). Overall, this network analysis suggested that migration and endothelial barrier restoration were strongly associated with genes regulating the transition of neural crest-derived cells to perivascular stromal cells (Subset 4 from SC2), and that, after migration, pleiotrophin may promote tight junction formation (restoring vascular integrity) rather than angiogenic new immature vessel formation.

## Discussion

In summary, using in vitro and in vivo experiments combined with gene profiling and interactome modeling, we have demonstrated that **(i)** a heretofore unrecognized repository of residual (“vestigial”) quiescent NCSCs persists into adulthood, sited in the neural crest-derived leptomeninges (and is a population distinct from previously-described neurogenic progenitors that migrated there from the neural tube (*Bifari et al., 2017; Decimo et al., 2011; Siegenthaler et al., 2009));* **(ii)** these NCSCs appear to perform a function in the adult (especially in response to a vascular injury like stroke) that is related to their role during embryogenesis: migrating to the perivascular space and restoring vascular integrity, in this case, BBB function. In this study, we focused on CD271+ cells derived from the leptomeninges. However, consistent with -- indeed, reinforcing -- our conclusion that these cells are residual NCSCs, CD271+ cells are also present in cranial sutures and skull bone, other embryonic neural crest derivatives, and may also be engaged for repair (Cai et al., 2019).

While we identified pleiotrophin as the candidate factor secreted by the leptomeningeal NCSCs for restoring BBB function, this factor can also be produced by other cells, such as pericytes (Nikolakopoulou et al., 2019). Therefore, future studies are warranted to investigate how other cell types interact with the leptomeningeal signaling mechanism described here.

Although we focused on stroke in this report as the prototypical vascular disruptor in the brain, perturbations in brain vasculature play critical roles in the pathophysiology of other neurological conditions, for example, in such neurodegenerative conditions as Alzheimer disease, Prion disease, and Parkinson disease (Da Mesquita et al., 2018; Sweeney et al., 2018). Future studies should explore whether residual NCSCs in the adult may be playing a role – or failing to do so -- in preserving endothelial barrier function in these vascular-related CNS disorders (**Fig. S6**). Our single-nucleus RNA-seq results provide a powerful database for aiding future investigations into the complex heterogeneity of perivascular subsets in the brain.

The neural crest has traditionally been regarded as a transient prenatal structure critical to embryogenesis (including mediating initial neurovascular co-patterning). Whether the neural crest – or remnants thereof -- genuinely persists (in a non-neoplastic state) beyond development – and certainly whether it performs any physiological function in the adult – has been less well understood and usually doubted. We provide evidence here that the meninges may be a repository of residual quiescent NCSCs in the adult mammalian brain, and that such cells may play a homeostasis-preserving role following CNS injury that is reminiscent of some of their actions in the embryo – preserving or restoring vascular integrity. This finding may also provide a mechanistic explanation for older observations that the meninges may be involved in the brain’s injury response (Bifari et al., 2017; Chen et al., 2019; Decimo et al., 2011; Decimo et al., 2012; Siegenthaler et al., 2009). Further studies are warranted to determine how this leptomeningeal mechanism and its mediators (e.g., pleiotrophin) may be exploited for developing new therapies against neurovascular dysfunction in CNS disease.

## Acknowledgments

We acknowledge funding from the Rappaport Foundation and NIH NINDS (R01NS115968). We thank Dr. Aviv Regev, Head Genentech Research and Early Development for discussions to coordinate experimental designs and support data analysis of snRNA-seq. We thank the Department of Pathology Flow and Image Cytometry Core at MGH149 for discussions to optimize cell gating. M.T. is supported by the International Centre for Translational Eye Research (MAB/2019/12) project which is carried out within the International Research Agendas programme of the Foundation for Polish Science co-financed by the European Union under the European Regional Development Fund. We thank B. Banasik for help with scVelo.

## Author Contributions

Performed experiments and/or analyzed data (YN, TN, JHP, MT^1^, EE, BJA, WL, VDL, RD, IS, ES); designed experiments (SS, EHL, ES, MT^5^, KH); wrote manuscript (EHL, ES, MT^5^, KH); funding and support (EHL, KH).

## SUPPLEMENTAL FIGURES

**Fig. S1:**
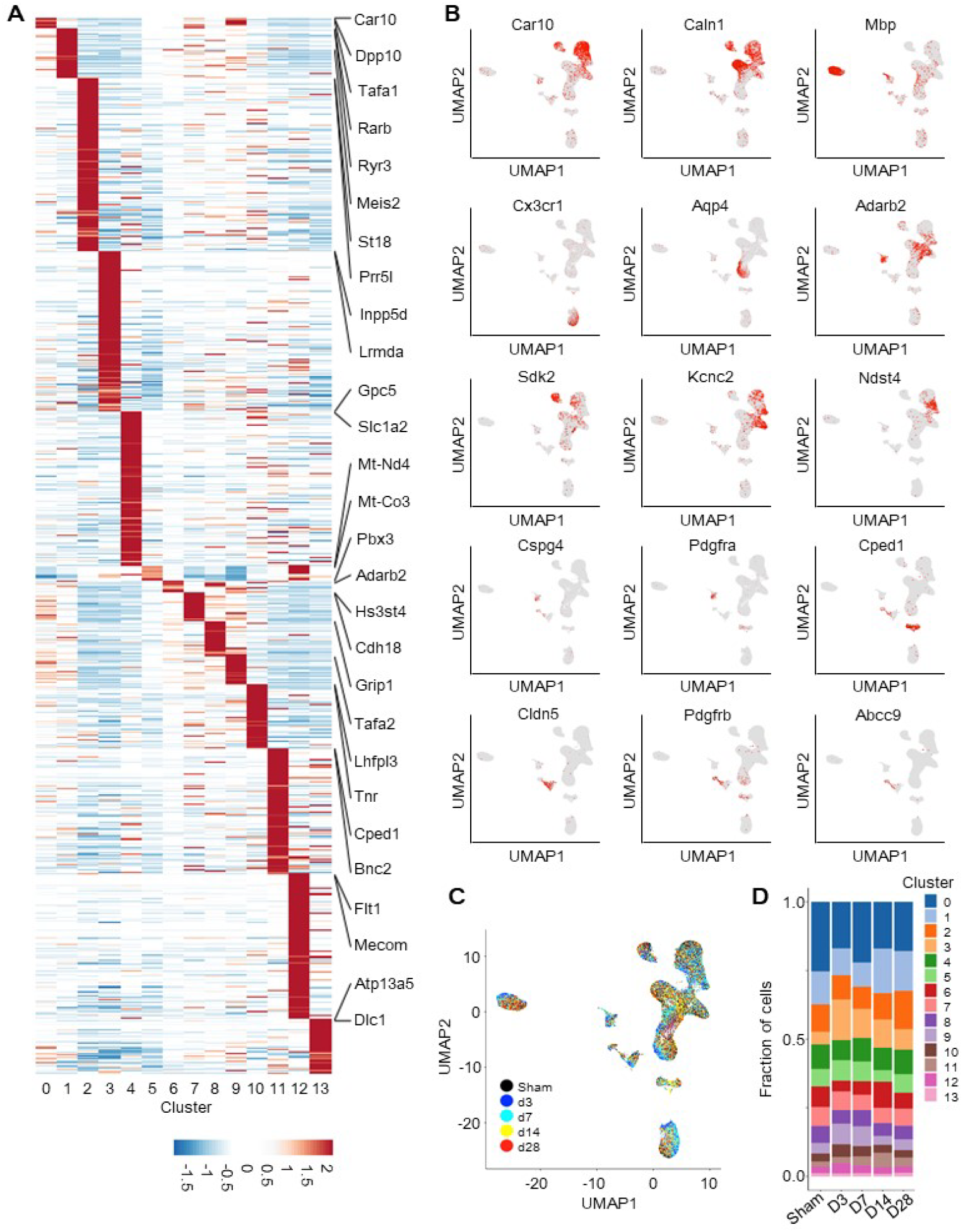
Differentially expressed gene (DEG) analysis. **A.** DEGs clustering revealed 0 to 13 clusters following single-nucleus RNA-seq. **B.** Representative markers for each cluster. **C.** Time course plots in UMAP. **D.** Census plots for each cluster.

**Fig. S2:**
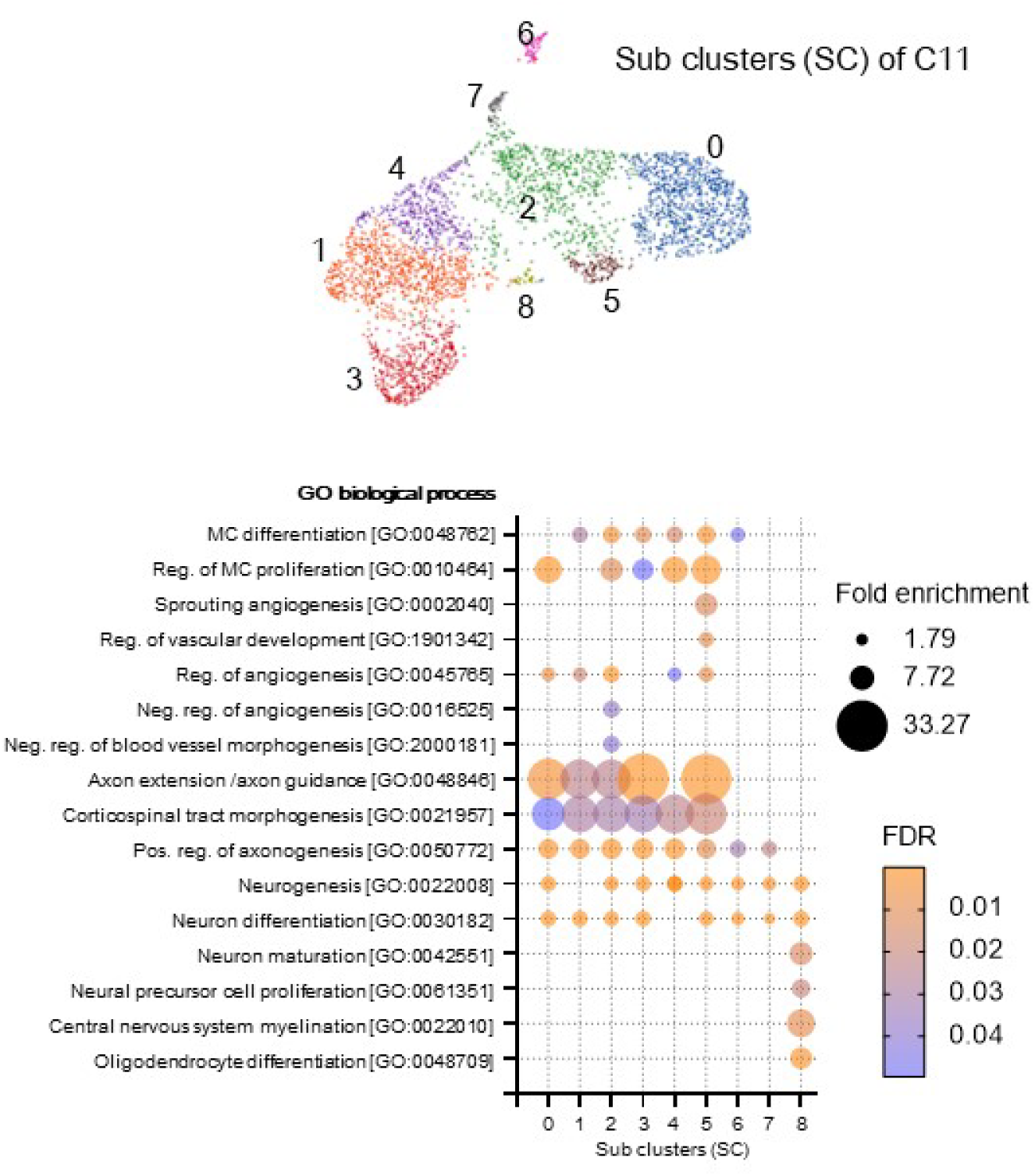
GO enrichment analysis in subclusters of Cluster 11: SC2, SC3 and SC5 commonly shared gene enrichment profile including the regulation of neural crest-derived mesenchymal differentiation, and proliferation, axon extension, and differentiation of neural crest-derived neurons. SC2 and SC5 also showed gene enrichment in the regulation of blood vessel morphogenesis while SC2 also upregulated genes involved in negative regulation of angiogenesis. Interestingly, SC8 uniquely upregulated genes regulating neuron maturation, neuronal precursor cell proliferation, and myelination.

**Fig. S3:**
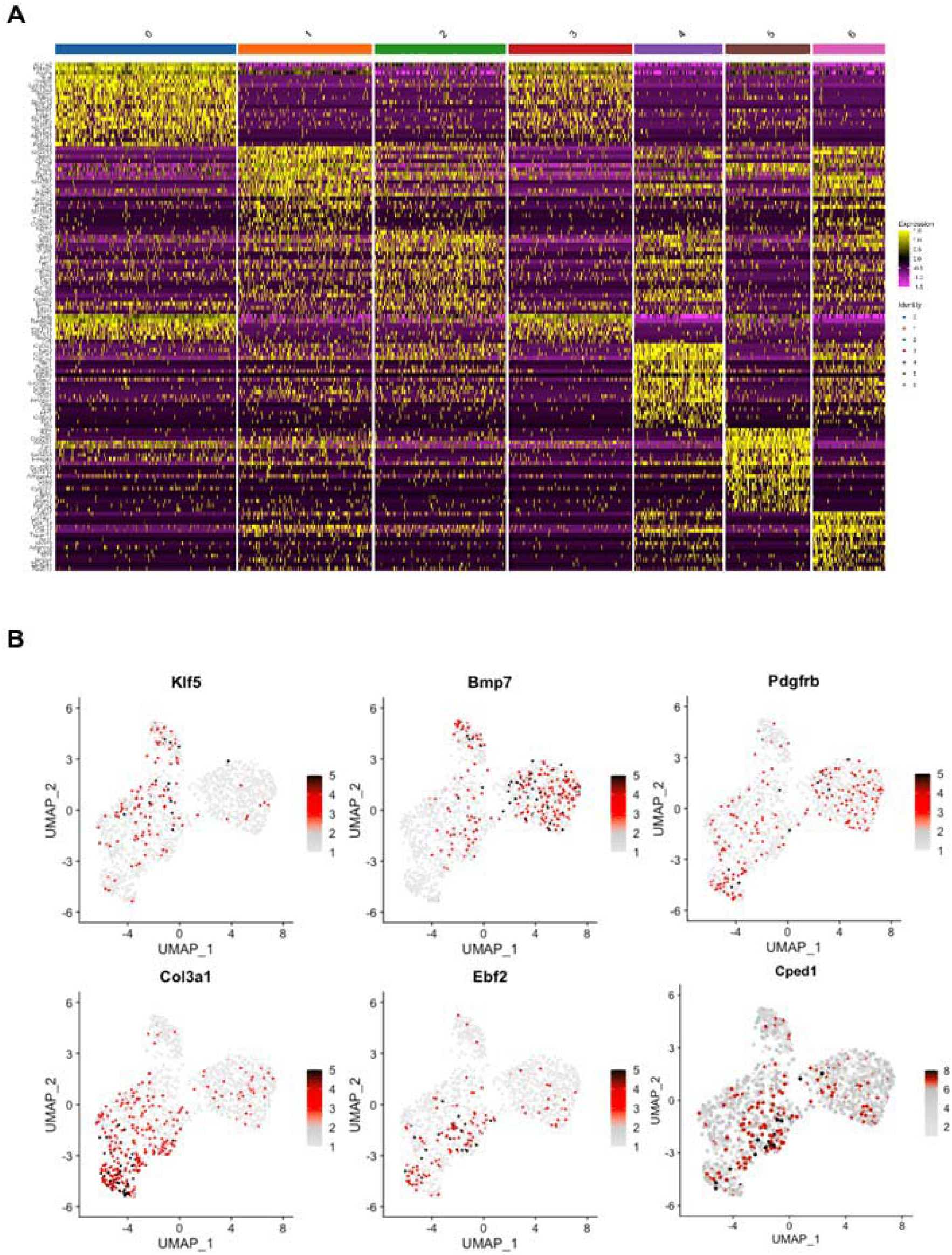
Deferential gene expression (DEG) in each subset – (A) heat map, (B) plots for top genes.

**Fig. S4:**
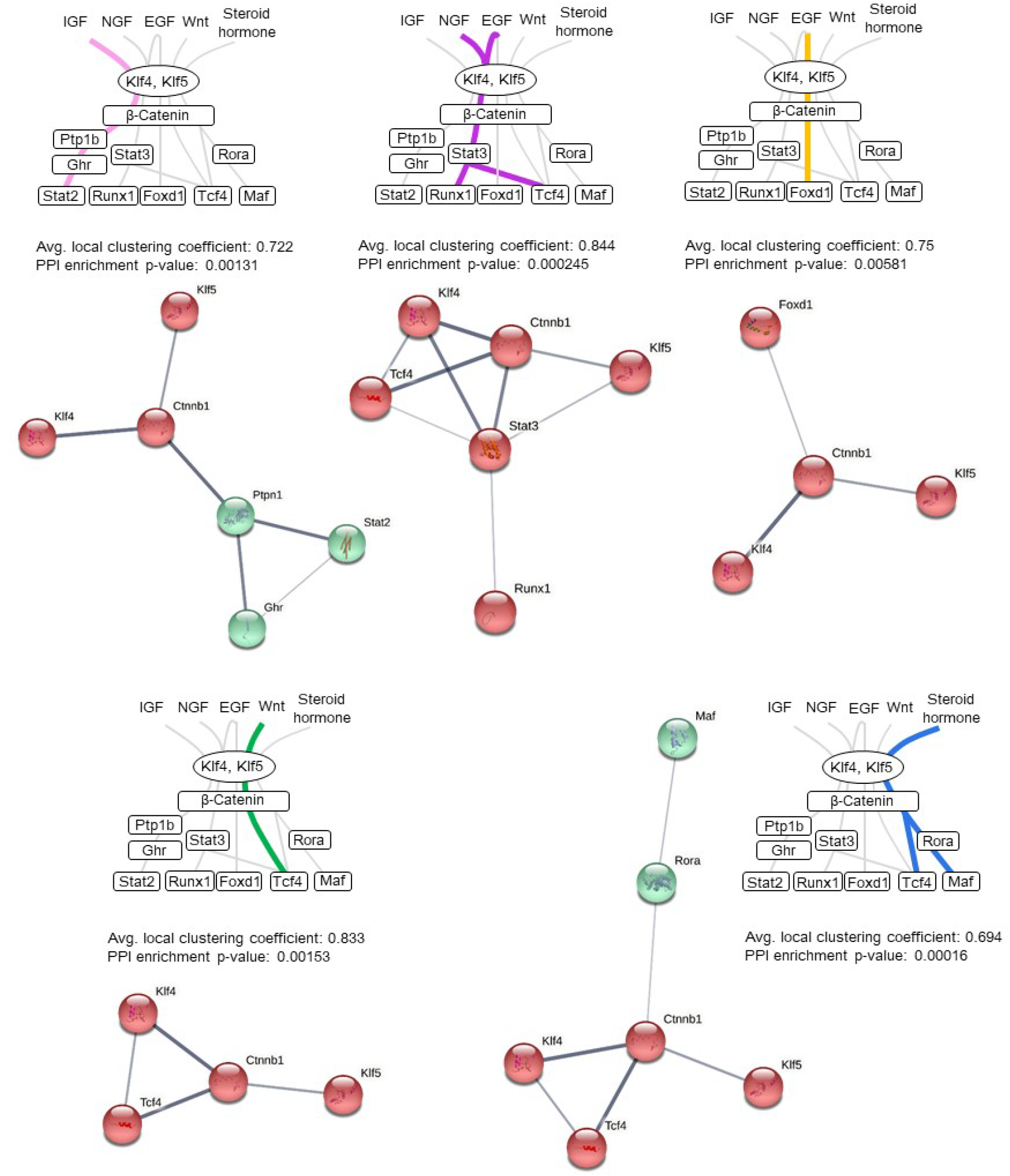
Protein-protein interaction (PPI) network in a transition molecular model analyzed by functional protein association networks: STRING. Each signaling pathway in this hypothetical molecular model had PPI enrichment p-values <0.05. All genes were clustered based on Markov cluster algorithm (MCL).

**Fig. S5:**
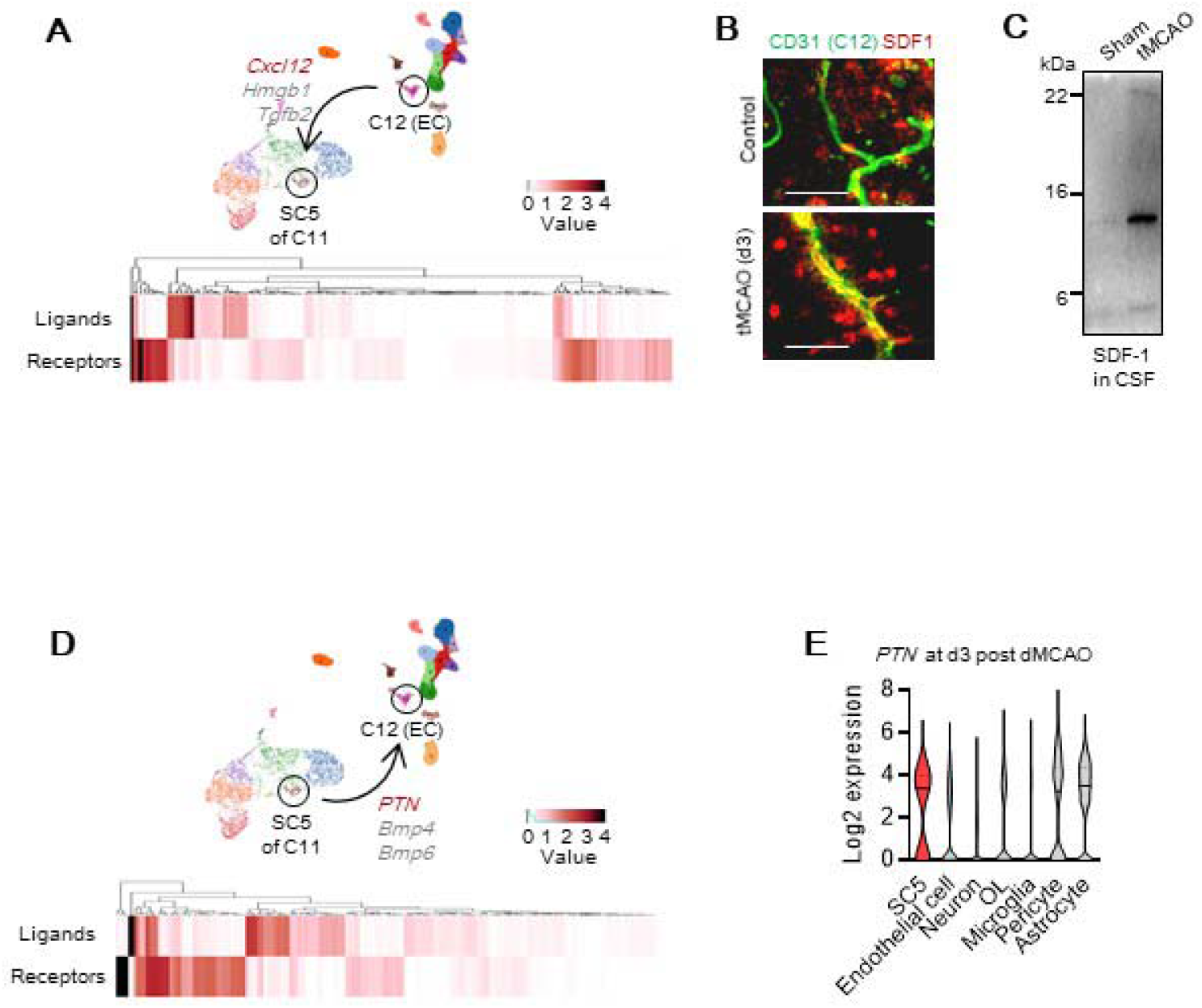
Receptor-ligand interactions. **A.** Receptor-ligand interaction analysis between SC5 and vascular endothelial cell (VEC) cluster C12 revealed endothelial *Cxcl12* (SDF-1α) as the highest expression value as ligand to SC5 subset. **B-C.** Immunostaining (**B**) and western blot (**C**) verified that SDF-1 α was upregulated in brain VECs and secreted into the cerebrospinal fluid (CSF) at 3 days after transient (60 min) focal cerebral ischemia in adult mice. **D.** Receptor-ligand interaction analysis between VEC cluster C12 and SC5 revealed pleiotrophin (PTN) as the highest expression value as ligand to the C12 VEC cluster. **E.** SC5 subset highly expressed PTN compared to other brain cell types

**Fig. S6:**
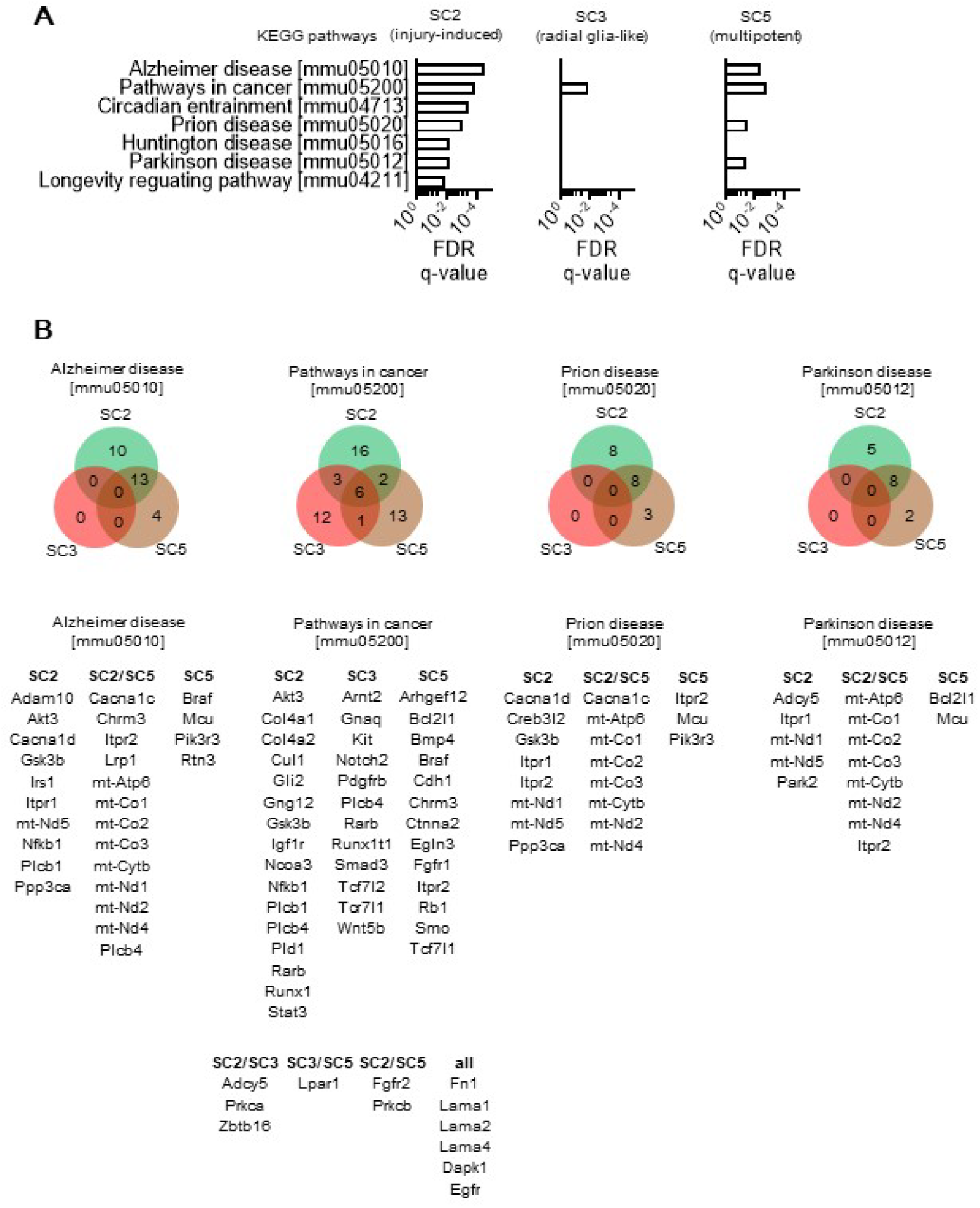
**A.** KEGG pathway analysis showed that SC2 and SC5 commonly enriched genes are involved in CNS vascular disorders other than solely stroke, e.g., Alzheimer disease, Pathways in Cancer, Prion disease, and Parkinson disease. **B.** Venn diagram shows common and unique genes expressed in SC2, SC3 or SC5 in each pathway.

## Materials and methods

### Chemicals

Brain-derived neurotrophic factor (BDNF, 450-02, Peprotech), beta-nerve growth factor (NGF, 450-01, Peprotech), Fibroblast growth factor (FGF, 100-18B, Peprotech), Insulin-like growth factor (IGF, 100-11B, Peprotech), recombinant human Pleiotrophin (252-PL-050, R&D systems), pAAV-CAG-tdTomato (#59462-AAV5, addgene), NeuroTrace^TM^ 640/660 Deep-Red (N21483, Thermo Fisher Scientific), FITC-dextran (FD40S, Sigma-Aldrich), recombinant human Pleiotrophin (252-PL-50, R&D systems).

### Antibodies

anti-CD271 antibodies (1:100, MA5-13311, Thermo Fisher Scientific, 1:100, 839701, BioLegend), anti-Foxd3 antibody (1:200, LS-C356037-100, LSBio), anti-β-actin antibody (1:10,000, A5441, Millipore-Sigma), anti-PDGFR-β antibody (1:100, AF1042, R&D Systems), anti-smooth muscle actin antibody (1:500, ab5694, Abcam), anti-TFAP2A antibody (1:100, sc-12726, Santacruz), anti-Sox10 antibody (1:200, 14-5923-82, Thermo Fisher Scientific), anti-CD57 (HNK) antibody (1:200, ab199156, Abcam), anti-Klf5 antibody (1:100, TA811868, Origene), anti-Bmp7 antibody (1:100, AF354, R&D Systems), anti-Col3a1 antibody (1:100, NB600-594, Novus Biologicals), anti-Cped1 antibody (1:100, PA5-52903, Thermo Fisher Scientific), anti-CD31 antibody (1:50, 565629, BD Biosciences), anti-Collagen (Type-IV) antibody (1:200, 1340-01, Sourthern Biotech), anti-LYVE-1 antibody (1:500, NB100-725B, Novus Biologicals), anti-Vimentin antibody (1:200, ab92547, Abcam), anti-CD90 antibody (1:200, sc-9163, Santacruz), anti-CD200 antibody (1:500, LS-C34538, LSBio), anti-pAkt antibody (1:50, 9271S, Cell Signaling), anti-pGSK3β antibody (1:50, sc-373800, Santacruz), anti-Claudin-5 antibody (1:100, ab131259, Abcam), anti-ZO1 antibody (1:100, 33-9100, Thermo Fisher Scientific).

### Animals

Male Sprague-Dawley rats (320 to 340 g) and male C57BL/6J mice (24–27 g) were housed in pathogen-free facilities with 12h day and night cycles. All experiments were performed following an institutionally approved protocol in accordance with National Institutes of Health guidelines and with the United States Public Health Service’s Policy on Human Care and Use of Laboratory Animals and following Animals in Research: Reporting *In vivo* Experiments (ARRIVE) guidelines.

### Cell cultures

Leptomeningeal CD271 cell cultures were prepared from male C57BL/6J mice (12-14 weeks) or male SD rats (12-14 weeks) subjected to focal cerebral ischemia or from human meningeal cells (1400, ScienCell). Briefly, Biotin-conjugated CD271 antibody was mixed with CELLection Dynabeads (11533D, Thermo Fisher Scientific) coated with recombinant streptavidin (100 μl, 4×10^7^ beads) for 30 min. After removing the free floating antibody, beads were mixed with cell suspensions for 20 min at 4C°. Isolated CD271+ cells were plated onto Geltrex-coated 6 well plates and cultured with DMEM/F12 plus 1% N2, 2% B27, FGF2 (100 ng/ml), and IGF1 (100 ng/ml) for 7-10 days. FBS (5%) was added to media based on cell condition.

### Models of focal cerebral ischemia

All animals were anesthetized with isoflurane (1.5%) in 30%/70% oxygen/nitrous oxide. Transient focal ischemia was induced introducing a 6-0 (in mice) or 5-0 (in rats) surgical monofilament nylon suture (Doccol) from the external carotid artery into the internal carotid artery and advancing it to the branching point of the MCA. Cerebral blood flow was monitored, by continuous laser doppler flowmetry (LDF) (Perimed, North Royalton, OH, U.S.A.), in the area of the MCA to confirm adequate occlusion. Animals that did not have a significant reduction to less than 30% baseline LDF values during MCAO were excluded. After occluding the MCA for 60 minutes in mice and 100 minutes in rats, the monofilament suture was gently withdrawn in order to restore blood flow, and LDF values were recorded for 10 minutes after reperfusion. As for permanent distal MCAO, a small craniotomy was made over the trunk of the left MCA and above the rhinal fissure. Distal MCAO was performed by placing a small piece of cellulose paper soaked with a 10% solution of FeCl_3_ in saline just before its bifurcation between the frontal and parietal branches. The paper remains placed on the artery for 5 minutes and after 20 minutes artery occlusion is completed. Complete interruption of blood flow was confirmed under an operating microscope. These experimental conditions led to moderately sized cortical infarcts. Rectal temperature was monitored and maintained at 37°C±0.5°C with a thermostatically controlled a heating pad during surgery and a heating lamp for 4 hours after surgery.

### Single-nucleus RNA-seq

Single nucleus RNA sequencing was conducted by Singulomics Corporation (https://singulomics.com/, Bronx NY). In summary, frozen mouse brain tissue samples were homogenized and lysed with Triton X-100 in RNase-free water for nuclei isolation. The isolated nuclei were purified, centrifuged, and resuspended in PBS with BSA and RNase Inhibitor. The nuclei were diluted to 700 nuclei/ul and loaded to 10x Genomics Chromium Controller to encapsulate single nuclei into droplet emulsions following the manufacturer’s recommendations (Pleasanton, CA, United States). Library preparation was performed according to the instructions in the Chromium Next GEM 3’ Single Cell Reagent kits v3.1. Amplified cDNAs and the libraries were measured by Qubit dsDNA HS assay (Thermo Fisher Scientific, Wilmington, DE) and quality assessed by BioAnalyzer (Agilent Technologies, Santa Clara, CA). Libraries were sequenced on a NovaSeq 6000 instrument (Illumina, San Diego, CA, United States), and reads were subsequently processed using 10x Genomics Cell Ranger analytical pipeline (v4.0) and mouse mm10 pre-mRNA reference.

### Single-Nucleus RNA-seq Library Preparation

Library preparation was performed according to the instructions in the Chromium Next GEM 3’ Single Cell Reagent kits v3.1. Amplified cDNAs and the libraries were measured by Qubit dsDNA HS assay (Thermo Fisher Scientific, Wilmington, DE) and quality assessed by BioAnalyzer (Agilent Technologies, Santa Clara, CA). Libraries were sequenced on a NovaSeq 6000 instrument (Illumina, San Diego, CA, United States) to an average depth of 135 million paired end reads per sample, with 150bp on the first read and 150bp on the second read.

### Single-Nucleus RNA Sequencing Data Analysis

Reads were processed using 10x Genomics Cell Ranger analytical pipeline (v4.0) and mouse mm10 pre-mRNA reference (10x Genomics). Dataset aggregation was performed using the Cellranger aggr function. Downstream analysis was performed using the R software package Seurat (v3.2.2, http://satijalab.org/seurat/) (Butler et al., 2018). We removed nucleus doublets by Scrublet (v0.2.1) (Wolock et al., 2019) and low-quality nuclei with less than 200 genes detected and removed genes expressed in fewer than 50 cells, leaving 107,020 cells and 20,785 genes for further analysis. The filtered raw gene expression matrix was cell-normalized over total number of counts, multiplied by 10,000 and log transformed. Principal Component Analysis (PCA) was performed on a submatrix of the top 1,000 most variable genes (computed using FindVariableGenes from Seurat). We evaluated the number of top principal components by the elbow method keeping 16 PCs for clustering and data visualization. The cells were clustered using a shared nearest neighbor (SNN) modularity optimization-based clustering algorithm (*FindClusters* in the Seurat package). To visualize cells in two dimensions, we used Uniform Manifold Approximation and Projection (UMAP) (Becht et al., 2018). The cluster-specific genes were computed using *FindAllMarkers* from the Seurat package, using the MAST test with the number of UMIs detected as a latent variable (Finak et al., 2015). The cluster-specific genes were computed using *FindAllMarkers* from the Seurat package, using the MAST test with the number of UMIs detected as a latent variable (Finak et al., 2015). Nucleus clusters were annotated by assessing known cell-type-specific markers (Batiuk et al., 2020; Rivera et al., 2021; Tasic et al., 2018; Zeisel et al., 2018) and by comparison with public mousebrain.org (http://mousebrain.org/adolescent/) and mouse brain and lung vascular and vessel-associated cell types (http://betsholtzlab.org/VascularSingleCells/database.html?gene=adgr14) datasets. Finegrained analysis of specific clusters was performed using the same methods and parameters except the number of top PCs that where adjusted accordingly.

### Creating gene signatures

Curated gene signatures were constructed from various databases of gene signatures as summarized in **Supplementary Table 3**. For a given nucleus, the expression values for each gene in the set were z-scored. Then, the signature of the nucleus has been determined as the mean z-score over all genes in the gene set.

### Determining potential receptor-ligand interactions

To identify potential paracrine signaling between cell-types, we leveraged method proposed by Schiebinger et al. First, we defined a list of ligands from the following GO terms: *cytokine activity* (GO:0005125), *growth factor activity* (GO:0008083), and *hormone activity* (GO:0005179). The set of receptors was defined by the GO term *receptor activity* (GO:0004872). Secondly, the expressed ligands and receptors in clusters of interest were intersected with the curated database of mouse protein-protein interactions. Finally, the average ligand and receptor expression values were computed within clusters of interest and the values were visually presented as heatmaps.

### Determining potential cellular trajectories

We leveraged RNA velocity method which uses spliced and unspliced forms of RNA transcripts to infer potential cellular trajectories. The bam files from 10x Genomics Cell Ranger pipeline were processed using Velocyto package (v.0.17) to determine spliced and unspliced RNA expression matrices. Then, the matrices were preprocessed using Scanpy toolkit (v1.7.2) and transition cell-cell transition probabilities were computed using scVelo (v0.2.3) with default parameters. Finally, we leveraged PAGA graph abstraction with velocity-inferred directionality to determine transitions between cell clusters and then PAGA velocity-graphs were visualized on the UMAP embedding.

Other analysis including GO term analysis (http://geneontology.org/), GSEA analysis (http://www.gsea-msigdb.org/gsea/msigdb/annotate.jsp) and KEGG pathway analysis (https://string-db.org/) were performed via each database.

### In vivo two photon imaging

Two weeks prior to in vivo two photon imaging, A total of 5×10^9^ viral genomes of pAAV-CAG-tdTomato (#59462-AAV5, addgene) was applied onto the cortical surface of a male C57BL/6J mouse (12-14 wks) through the cranial window. Before and d3 post-stroke, fluorescein-conjugated dextran was infused intravenously to visualize blood vessels and the cranial window with a glass coverslip was illuminated with a laser 840 nm to capture tdTomato+ cells.

### Blood-Brain Barrier (BBB) permeability

BBB permeability of mice was evaluated 4 days after MCAO by Evans blue dye. Briefly, 2% Evans blue dye (E2129-10G, Sigma) in 0.9% saline (4 mL/kg) was injected intravenously into the tail vein and was allowed to circulate for 2 hours. Removal of intravascular dyes was performed by perfusion of phosphate-buffered saline (PBS) via left cardiac ventricle in mice. For quantitative measurement of Evans blue leakage, the ipsilateral and contralateral hemisphere was removed, weighted and homogenized in 50% trichloroacetic acid solution, then centrifuged at 15,000 rpm for 30 min. Evans blue concentration was quantitatively determined by measuring the 610 nm absorbance of supernatant.

### Western blot analysis

Cultures were rinsed twice with PBS and cell membrane extracts were collected according to the method of ProteoExtract Kit (Calbiochem). Samples were heated with equal volumes of SDS sample buffer (Novex) and 10% 2-mercaptoethanol at 95°C for 5 min, and then each sample was loaded onto 4-20% Tris-glycine gels. After electrophoresis and transferred to polyvinylidene difluoride membranes (Novex), the membranes were blocked in Tris-buffered saline containing 0.1% Tween 20 and 0.2% I-block (Tropix) for 60 min at 4°C. Membranes were then incubated overnight at 4°C with primary antibodies. After incubation with peroxidase-conjugated secondary antibodies, chemiluminescence was enhanced (GE Healthcare). Optical density was assessed using the NIH Image analysis software.

### Immunohistochemistry

Cells were washed with ice-cold PBS (pH 7.4), followed by 4% paraformaldehyde for 10 min. After being washed three times in PBS containing 0.1% Triton X-100, they were incubated with 3% bovine serum albumin (BSA) in PBS for 1 h. Then cells were incubated with primary antibodies at 4°C overnight. After washing with PBS, they were incubated with secondary antibodies (1:400; Jackson ImmunoResearch) for 1 h at room temperature. Finally, nuclei were counterstained with 4,6-diamidino-2-phenylindole (DAPI), and coverslips were placed. Immunostaining was analyzed with a fluorescence microscope (Nikon ECLIPSE Ti-S).

### Quantitative PCR analysis

Total RNAs were extracted using QIAzol lysis reagent (QIAGEN, 79306) from isolated CD271 cells, followed by cDNA synthesis using High-capacity RNA-to-cDNA kit (ThermoFisher Seicntific, 4387406) according to the manufacturer’s instructions. Relative levels of genes were determined by amplifying FOXD3 (Applied Biosystems, Mm02384867_s1), TFAP2 (Applied Biosystems, Mm00493468_m1), SOX10 (Applied Biosystems, Mm00569909_m1), NGFR (Applied Biosystems, Mm00446296_m1), PRRX2 (Applied Biosystems, Mm00436428_m1), TWIST1 (Applied Biosystems, Mm00442036_m1) and normalized by housekeeping gene HPRT (Applied Biosystems, Mm01545399_m1).

### FACS analysis

FACS analysis was performed as described before (Hayakawa et al., 2016; Hayakawa et al., 2012). Tissues collected from leptomeninges and peri-infarct cortex are gently minced and then digested at 37°C for 30 min with an enzyme cocktail (Collagenase type IV; Sigma-Aldrich, C5138, DNase I; Sigma-Aldrich, D4263). FACS analysis was performed to determine tdTomato+ and CD271 FITC using BD Fortessa with a no labeled control for determining appropriate gates, voltages, and compensations required in multivariate flow cytometry. Data was analyzed by FlowJo or Flowing Software (https://bioscience.fi/services/cell-imaging/flowing-software/).

### Statistical analysis

All of the experiments were performed in duplicate, repeated 3-5 times independently. All of in vivo experiments were performed with full blinding, allocation concealment and randomization. GraphPad Prism version 9 was used overall statistical analysis in this study. Results were expressed as mean±SD. When only two groups were compared, the Student’s t-test was used. Multiple comparisons were evaluated by Tukey’s test after one-way ANOVA. *P*<0.05 was considered to be statistically significant.

## References

Acevedo, L.M., Lindquist, J.N., Walsh, B.M., Sia, P., Cimadamore, F., Chen, C., Denzel, M., Pernia, C.D., Ranscht, B., Terskikh, A., et al. (2015). hESC Differentiation toward an Autonomic Neuronal Cell Fate Depends on Distinct Cues from the Co-Patterning Vasculature. Stem Cell Reports 4, 1075–1088.

Barque, A., Jan, K., De La Fuente, E., Nicholas, C.L., Hynes, R.O., and Naba, A. (2021). Knockout of the gene encoding the extracellular matrix protein SNED1 results in early neonatal lethality and craniofacial malformations. Dev Dyn 250, 274–294.

Batarfi, M., Valasek, P., Krejci, E., Huang, R. and Patel, K. (2017). The development and origins of vertebrate meninges. Biological Communications 62, 73–81.

Bifari, F., Decimo, I., Pino, A., Llorens-Bobadilla, E., Zhao, S., Lange, C., Panuccio, G., Boeckx, B., Thienpont, B., Vinckier, S., et al. (2017). Neurogenic Radial Glia-like Cells in Meninges Migrate and Differentiate into Functionally Integrated Neurons in the Neonatal Cortex. Cell Stem Cell 20, 360–373 e367.

Brundage, M.E., Tandon, P., Eaves, D.W., Williams, J.P., Miller, S.J., Hennigan, R.H., Jegga, A., Cripe, T.P., and Ratner, N. (2014). MAF mediates crosstalk between Ras-MAPK and mTOR signaling in NF1. Oncogene 33, 5626–5636.

Cai, R., Pan, C., Ghasemigharagoz, A., Todorov, M.I., Forstera, B., Zhao, S., Bhatia, H.S., Parra-Damas, A., Mrowka, L., Theodorou, D., et al. (2019). Panoptic imaging of transparent mice reveals whole-body neuronal projections and skull-meninges connections. Nat Neurosci 22, 317–327.

Chen, J., He, J., Ni, R., Yang, Q., Zhang, Y., and Luo, L. (2019). Cerebrovascular Injuries Induce Lymphatic Invasion into Brain Parenchyma to Guide Vascular Regeneration in Zebrafish. Dev Cell 49, 697–710 e695.

Da Mesquita, S., Louveau, A., Vaccari, A., Smirnov, I., Cornelison, R.C., Kingsmore, K.M., Contarino, C., Onengut-Gumuscu, S., Farber, E., Raper, D., et al. (2018). Functional aspects of meningeal lymphatics in ageing and Alzheimer’s disease. Nature 560, 185–191.

Decimo, I., Bifari, F., Rodríguez, F.J., Malpeli, G., Dolci, S., Lavarini, V., Pretto, S., Vasquez, S., Sciancalepore, M., Montalbano, A., et al. (2011). Nestin- and doublecortin-positive cells reside in adult spinal cord meninges and participate in injury-induced parenchymal reaction. Stem Cells 29, 2062–2076.

Decimo, I., Dolci, S., Panuccio, G., Riva, M., Fumagalli, G., and Bifari, F. (2020). Meninges: A Widespread Niche of Neural Progenitors for the Brain. Neuroscientist, 1073858420954826.

Decimo, I., Fumagalli, G., Berton, V., Krampera, M., and Bifari, F. (2012). Meninges: from protective membrane to stem cell niche. Am J Stem Cells 1, 92–105.

Dupin, E., and Sommer, L. (2012). Neural crest progenitors and stem cells: from early development to adulthood. Dev Biol 366, 83–95.

Etchevers, H.C., Vincent, C., Le Douarin, N.M., and Couly, G.F. (2001). The cephalic neural crest provides pericytes and smooth muscle cells to all blood vessels of the face and forebrain. Development 128, 1059–1068.

Garcez, R.C., Teixeira, B.L., Schmitt Sdos, S., Alvarez-Silva, M., and Trentin, A.G. (2009). Epidermal growth factor (EGF) promotes the in vitro differentiation of neural crest cells to neurons and melanocytes. Cell Mol Neurobiol 29, 1087–1091.

Hallmann, R., Horn, N., Selg, M., Wendler, O., Pausch, F., and Sorokin, L.M. (2005). Expression and function of laminins in the embryonic and mature vasculature. Physiol Rev 85, 979–1000.

Hari, L., Brault, V., Kleber, M., Lee, H.Y., llle, F., Leimeroth, R., Paratore, C., Suter, U., Kemler, R., and Sommer, L. (2002). Lineage-specific requirements of beta-catenin in neural crest development. J Cell Biol 159, 867–880.

La Manno, G., Soldatov, R., Zeisel, A., Braun, E., Hochgerner, H., Petukhov, V., Lidschreiber, K., Kastriti, M.E., Lonnerberg, P., Furlan, A., et al. (2018). RNA velocity of single cells. Nature 560, 494–498.

Lee, G., Kim, H., Elkabetz, Y., Al Shamy, G., Panagiotakos, G., Barberi, T., Tabar, V., and Studer, L. (2007). Isolation and directed differentiation of neural crest stem cells derived from human embryonic stem cells. Nat Biotechnol 25, 1468–1475.

Nakagomi, T., Molnar, Z., Nakano-Doi, A., Taguchi, A., Saino, O., Kubo, S., Clausen, M., Yoshikawa, H., Nakagomi, N., and Matsuyama, T. (2011). Ischemia-induced neural stem/progenitor cells in the pia mater following cortical infarction. Stem Cells Dev 20, 2037–2051.

Nikolakopoulou, A.M., Montagne, A., Kisler, K., Dai, Z., Wang, Y., Huuskonen, M.T., Sagare, A.P., Lazic, D., Sweeney, M.D., Kong, P., et al. (2019). Pericyte loss leads to circulatory failure and pleiotrophin depletion causing neuron loss. Nat Neurosci 22, 1089–1098.

Olesnicky Killian, E.C., Birkholz, D.A., and Artinger, K.B. (2009). A role for chemokine signaling in neural crest cell migration and craniofacial development. Dev Biol 333, 161–172.

Siegenthaler, J.A., Ashique, A.M., Zarbalis, K., Patterson, K.P., Hecht, J.H., Kane, M.A., Folias, A.E., Choe, Y., May, S.R., Kume, T., et al. (2009). Retinoic acid from the meninges regulates cortical neuron generation. Cell 139, 597–609.

Simoes-Costa, M., and Bronner, M.E. (2016). Reprogramming of avian neural crest axial identity and cell fate. Science 352, 1570–1573.

Soldatov, R., Kaucka, M., Kastriti, M.E., Petersen, J., Chontorotzea, T., Englmaier, L., Akkuratova, N., Yang, Y., Haring, M., Dyachuk, V., et al. (2019). Spatiotemporal structure of cell fate decisions in murine neural crest. Science 364.

Sweeney, M.D., Kisler, K., Montagne, A., Toga, A.W., and Zlokovic, B.V. (2018). The role of brain vasculature in neurodegenerative disorders. Nat Neurosci 21, 1318–1331.

Vanlandewijck, M., He, L., Mae, M.A., Andrae, J., Ando, K., Del Gaudio, F., Nahar, K., Lebouvier, T., Lavina, B., Gouveia, L., et al. (2018). A molecular atlas of cell types and zonation in the brain vasculature. Nature 554, 475–480.

Weigele, J., and Bohnsack, B.L. (2020). Genetics Underlying the Interactions between Neural Crest Cells and Eye Development. J Dev Biol 8.

Williams, R.M., Candido-Ferreira, I., Repapi, E., Gavriouchkina, D., Senanayake, U., Ling, I.T.C., Telenius, J., Taylor, S., Hughes, J., and Sauka-Spengler, T. (2019). Reconstruction of the Global Neural Crest Gene Regulatory Network In Vivo. Dev Cell 51, 255–276 e257.

Wolf, F.A., Harney, F.K., Plass, M., Solana, J., Dahlin, J.S., Gottgens, B., Rajewsky, N., Simon, L., and Theis, F.J. (2019). PAGA: graph abstraction reconciles clustering with trajectory inference through a topology preserving map of single cells. Genome Biol 20, 59.

Yang, Y., and Torbey, M.T. (2020). Angiogenesis and Blood-Brain Barrier Permeability in Vascular Remodeling after Stroke. Curr Neuropharmacol 18, 1250–1265.

## References for Methods and materials

Batiuk, M.Y., Martirosyan, A., Wahis, J., de Vin, F., Marneffe, C., Kusserow, C., Koeppen, J., Viana, J.F., Oliveira, J.F., Voet, T., et al. (2020). Identification of region-specific astrocyte subtypes at single cell resolution. Nat Commun 11, 1220.

Becht, E., McInnes, L., Healy, J., Dutertre, C.A., Kwok, I.W.H., Ng, L.G., Ginhoux, F., and Newell, E.W. (2018). Dimensionality reduction for visualizing single-cell data using UMAP. Nat Biotechnol.

Butler, A., Hoffman, P., Smibert, P., Papalexi, E., and Satija, R. (2018). Integrating single-cell transcriptomic data across different conditions, technologies, and species. Nat Biotechnol 36, 411–420.

Finak, G., McDavid, A., Yajima, M., Deng, J., Gersuk, V., Shalek, A.K., Slichter, C.K., Miller, H.W., McElrath, M.J., Prlic, M., et al. (2015). MAST: a flexible statistical framework for assessing transcriptional changes and characterizing heterogeneity in single-cell RNA sequencing data. Genome Biol 16, 278.

Hayakawa, K., Esposito, E., Wang, X., Terasaki, Y., Liu, Y., Xing, C., Ji, X., and Lo, E.H. (2016). Transfer of mitochondria from astrocytes to neurons after stroke. Nature 535, 551–555.

Hayakawa, K., Pham, L.D., Katusic, Z.S., Arai, K., and Lo, E.H. (2012). Astrocytic high-mobility group box 1 promotes endothelial progenitor cell-mediated neurovascular remodeling during stroke recovery. Proc Natl Acad Sci U S A 109, 7505–7510.

Rivera, A.D., Pieropan, F., Chacon-De-La-Rocha, I., Lecca, D., Abbracchio, M.P., Azim, K., and Butt, A.M. (2021). Functional genomic analyses highlight a shift in Gpr17-regulated cellular processes in oligodendrocyte progenitor cells and underlying myelin dysregulation in the aged mouse cerebrum. Aging Cell 20, e13335.

Tasic, B., Yao, Z., Graybuck, L.T., Smith, K.A., Nguyen, T.N., Bertagnolli, D., Goldy, J., Garren, E., Economo, M.N., Viswanathan, S., et al. (2018). Shared and distinct transcriptomic cell types across neocortical areas. Nature 563, 72–78.

Wolock, S.L., Lopez, R., and Klein, A.M. (2019). Scrublet: Computational Identification of Cell Doublets in Single-Cell Transcriptomic Data. Cell Syst 8, 281–291 e289.

Zeisel, A., Hochgerner, H., Lonnerberg, P., Johnsson, A., Memic, F., van der Zwan, J., Haring, M., Braun, E., Borm, L.E., La Manno, G., et al. (2018). Molecular Architecture of the Mouse Nervous System. Cell 174, 999–1014 e1022.

